# Formation of Multinucleated Osteoclasts Depends on an Oxidized Species of Cell Surface Associated La Protein

**DOI:** 10.1101/2024.05.02.592254

**Authors:** Evgenia Leikina, Jarred M. Whitlock, Kamran Melikov, Wendy Zhang, Michael P. Bachmann, Leonid V. Chernomordik

**Affiliations:** Section on Membrane Biology, Eunice Kennedy Shriver National Institute of Child Health and Human Development, National Institutes of Health, Bethesda, MD 20892, USA; Department of Radioimmunology, Institute of Radiopharmaceutical Cancer Research, Helmholtz-Zentrum Dresden-Rossendorf (HZDR), 01328 Dresden, Germany; Institute of Immunology, Medical Faculty Carl Gustav Carus Dresden, Technical University Dresden, 01307 Dresden, Germany; University Cancer Center (UCC), Tumor Immunology, University Hospital Carl Gustav Carus Dresden, Technical University Dresden, 01307 Dresden, Germany

## Abstract

The bone-resorbing activity of osteoclasts plays a critical role in the life-long remodeling of our bones that is perturbed in many bone loss diseases. Multinucleated osteoclasts are formed by the fusion of precursor cells, and larger cells - generated by an increased number of cell fusion events - have higher resorptive activity. We find that osteoclast fusion and bone-resorption are promoted by reactive oxygen species (ROS) signaling and by an unconventional low molecular weight species of La protein, located at the osteoclast surface. Here, we develop the hypothesis that La’s unique regulatory role in osteoclast multinucleation and function is controlled by a ROS switch in La trafficking. Using antibodies that recognize reduced or oxidized species of La, we find that differentiating osteoclasts enrich an oxidized species of La at the cell surface, which is distinct from the reduced La species conventionally localized within cell nuclei. ROS signaling triggers the shift from reduced to oxidized La species, its dephosphorylation and delivery to the surface of osteoclasts, where La promotes multinucleation and resorptive activity. Moreover, intracellular ROS signaling in differentiating osteoclasts oxidizes critical cysteine residues in the C-terminal half of La, producing this unconventional La species that promotes osteoclast fusion. Our findings suggest that redox signaling induces changes in the location and function of La and may represent a promising target for novel skeletal therapies.

## INTRODUCTION

The integrity of our bones throughout life depends on tightly regulated coordination between the bone-forming activity of osteoblasts and the bone-resorbing activity of osteoclasts ^1–3^. A variety of genetic and age-related skeletal disorders are linked to a disbalance in osteoblast-osteoclast functional coupling that commonly results in excessive bone resorption and/or insufficient synthesis and mineralization of bone.

Multinucleated osteoclasts are formed by the fusion of mononucleated precursor cells, and, in most cases, cells with more nuclei (i.e., generated by a larger number of fusion events) have higher resorptive activity ^4,5^. We recently demonstrated that both osteoclast fusion and bone-resorption are controlled by an unconventional, low molecular weight form of La protein ^6^. La (SSB small RNA binding exonuclease protection factor La, Gene ID 6741, NCBI GENE), an abundant, ubiquitous protein in eukaryotes, exists primarily as a phosphorylated, nuclear species that plays essential functions in the maturation of RNA polymerase III transcripts, particularly tRNA ^7,8^. However, we discovered that La is dephosphorylated, proteolytically cleaved, and delivered to the surface of osteoclasts during multinucleation. Surface-associated La promotes cell-cell fusion and increased resorptive capacity in osteoclasts. When osteoclasts reach a mature size appropriate for their biological activity, fusion stops. Completion of the fusion process coincides with the removal of surface-associated La, suggesting that the changes in the surface La amounts effectively set the size and biological activity of osteoclasts. The mechanisms that trigger the radical switch in the location and function of La from nucleus to cell surface; and from ubiquitous RNA chaperone to an osteoclast-specific fusion regulator - remain to be understood.

Intracellular ROS represent a common biological switch for proteins with multiple functions in eukaryotes ^9,10^. Excessive levels of ROS during oxidative stress eventually lead to irreversible damage of biological systems and tissues and have been linked to diverse diseases in many physiological systems, including the skeleton ^11^. However, transient, moderate increases in ROS levels, referred to as redox signaling ^12,13^ or a mild oxidative stress ^14^, play important roles in diverse cellular differentiation processes^15–17^. Intracellular ROS signaling commonly induces the formation of disulfide bonds, drives structural and oligomeric transitions, and promotes the unconventional secretion of some proteins lacking a signal peptide ^18–20^. Alternatively, a transition in the redox state of cysteine residues can also be triggered merely by protein trafficking changes that shifts the localization of a protein from the cytosol (typically reducing) to the extracellular environment (typically oxidizing) ^21,22^.

Like many other physiological processes, bone remodeling and, more specifically, osteoclast formation depend on ROS signaling ^17,23^. RANKL-induced differentiation of osteoclast precursors quickly generates transient ROS signaling in differentiating osteoclasts ^24^, and many bone diseases, including osteoporosis, have been linked to perturbations in ROS signaling ^17,25^. Moreover, application of oxidizing reagents, such as H_2_O_2_, promotes osteoclast formation ^26–30^. In contrast, cell-permeable antioxidants block RANKL-induced ROS production and inhibit osteoclast formation and bone resorption ^31–34^.

Recent biochemical studies demonstrate that oxidizing conditions and intracellular redox signaling elicit conformational transitions and oligomerization of La protein and promote its nucleus-to-cytoplasm shuttling ^35,36^. Here, we tested the hypothesis that La’s unique regulatory role in osteoclast multinucleation and function is controlled by a ROS switch in La trafficking and function. Using antibodies that recognize reduced vs oxidized species of La, we found that nuclear La and cell surface La in differentiating osteoclasts to represent reduced and oxidized species of the protein, respectively. Oxidized La species at the surface of osteoclasts promoted their fusion and increased multinucleation. Suppressing ROS signaling in osteoclast precursors inhibited the appearance of La in the cytoplasm and at the surface of osteoclasts and suppressed fusion during osteoclast formation. Addition of the C-terminal half of La – the region required for promoting osteoclast fusion ^6^ – to the extracellular surface of osteoclasts rescued this inhibition but only if critical La cysteine residues were available for oxidation. Our data suggest that transient ROS signaling induces a shift from reduced to oxidized La species and plays a critical role in directing the delivery of La to the surface of osteoclasts and promoting multinucleation and subsequent resorptive function of these syncytial bone remodelers. Our findings suggest that redox transition and, more specifically, La cysteine oxidation may represent promising targets for therapeutic strategies aimed at modulating osteoclast-dependent bone resorption in skeletal pathologies.

## RESULTS

### Osteoclast fusion depends on an oxidized form of surface La

To mechanistically evaluate the transition from osteoclast precursors to multinucleated osteoclasts, we incubated primary human monocytes with recombinant macrophage colony-stimulating factor (M-CSF) and then with M-CSF and recombinant receptor activator of NF-kappaB ligand (RANKL) ^6,37^. While the time course and efficiency of multinucleated osteoclast formation varies from donor to donor ^6^, on average, we observed the appearance of small, multinucleated syncytia two days after RANKL application and fusion rapidly increased over the next two days, resulting in mature, resorption competent osteoclasts at 4 to 5 days post-RANKL addition ^6^. Previously, we demonstrated that the cell fusion stage of multinucleated osteoclast formation depends on La trafficking to the surface of the osteoclasts at days 2 to 4 post RANKL application ^6^, however the mechanisms that trigger La’s trip to the osteoclast surface and what molecular requirements must be met for its unconventional fusion role there remained open questions.

To address open questions concerning La’s surface trafficking and molecular function in osteoclast multinucleation, we evaluated the redox state of La in fusing osteoclasts using recently validated monoclonal α-La antibodies that recognize oxidized La (clone 7B6) or reduced La (clone 312B), or do not distinguish between these La species (Pan, clone 5B9) ^35^. At the time of fusion, immunofluorescence analysis of La’s localization in permeabilized osteoclasts with pan α-La antibody demonstrated that La is present in both the cytoplasm and nuclei (Fig. 1a). In stark contrast, the cytoplasmic/plasma membrane associated pool was recognized by the α-La antibody that recognizes oxidized La species, while the α-La antibody that recognizes reduced La species recognized the nuclear protein pool (Fig. 1a; Supp. Fig. 1). Under non-permeabilizing conditions – focusing on the exofacial surface of the plasma membrane - we readily observed La at the surface of fusing osteoclasts (see also ^6^). We find that this surface pool of La is dramatically enriched in oxidized, rather than reduced, La species (Fig. 1b). In further support of this conclusion, we found that application of the membrane-impermeable reducing reagent Tris (2-carboxyethyl) phosphine (TCEP) dramatically decreased the recognition of surface La by the α-La antibody that recognizes the oxidized species and increased the recognition of surface La by the α-La antibody that recognizes the reduced species (Fig. 1b-c). In contrast, TCEP had no effect on the ability of the pan α-La antibody to recognize surface La (Fig. 1b-c) or on the surface detection of La with another α-La antibody (Abcam #75927) that we have used previously to evaluate La at the surface of fusing osteoclasts ^6^ and now found to recognize both reduced and oxidized species of La (data not shown). These findings indicate that the surface pool of La that manages osteoclast size and resorptive function is primarily composed of an oxidized La molecular species. Interestingly, TCEP treatment did not dissociate cell surface La from plasma membrane, suggesting that the reduced species of La retains its ability to associate with the plasma membrane.

**Figure 1:**
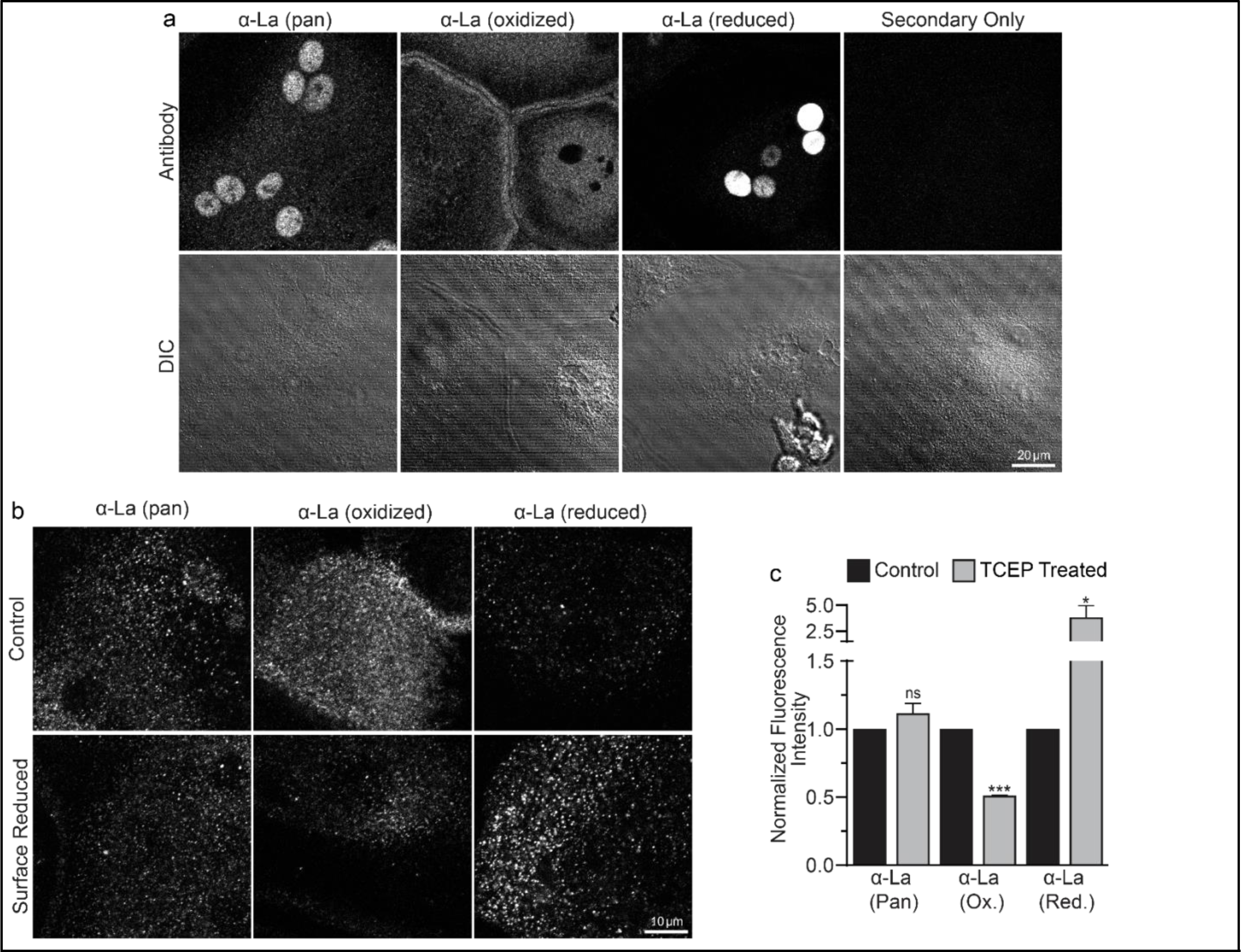
Oxidized La decorates the surface of osteoclasts during multinucleation. **(a)** Representative immunofluorescence (top) and differential interference contrast (DIC, bottom) confocal micrographs of permeabilized primary human osteoclasts. La localization was visualized via a general α-La antibody (Pan), an α-La antibody that recognizes oxidized La, or an α-La antibody thar recognizes reduced La (described and validated in {Berndt, 2021 #2}). **(b)** Representative immunofluorescence confocal micrographs of primary human osteoclasts stained with the antibodies described in **a** under non-permeabilized conditions to visualize surface La. Surface La pools were visualized for the untreated cells (control) and for cells treated with the membrane impermeable reducing reagent TCAP. **(c)** Quantification of **b**. (n=3) (p= 0.13, 0.0001, and 0.03, respectively. Statistical significance evaluated via paired t-test.

We further assessed the functional importance of surface La’s redox status in synchronized osteoclast fusion. As in earlier studies ^6,38,39^, we uncoupled the cell fusion stage of osteoclast formation from pre-fusion differentiation processes using lysophosphatidylcholine (LPC), a reversible inhibitor of an early stage of membrane rearrangement required for osteoclast fusion. LPC was applied for 16h following 2 days of RANKL elicited osteoclastogenesis. Ready-to-fuse cells, which could not fuse in the presence of LPC, rapidly fused after LPC wash out (Fig. 2a,b). As we demonstrated previously with another pan α-La antibody ^6^, pan α-La antibody 5B9 applied at the time of LPC removal inhibited synchronized osteoclast fusion (Fig. 2a,b). Similarly, α-La antibodies that recognize the oxidized species inhibited fusion, while antibodies that recognize the reduced La species had no effect on fusion (Fig. 2a,b). These data strongly support the conclusion that the functional La species that promotes osteoclast fusion and function is an unconventional, cell surface-associated, oxidized species. In further support of the functional importance of surface La oxidation, and possibly other surface proteins, we find that reducing the surface of human osteoclasts via TCEP treatment — which reduces surface La but does not alter its membrane association (Fig. 1c) — inhibits synchronized osteoclast fusion in a dose dependent manner (Fig. 2c).

**Figure 2:**
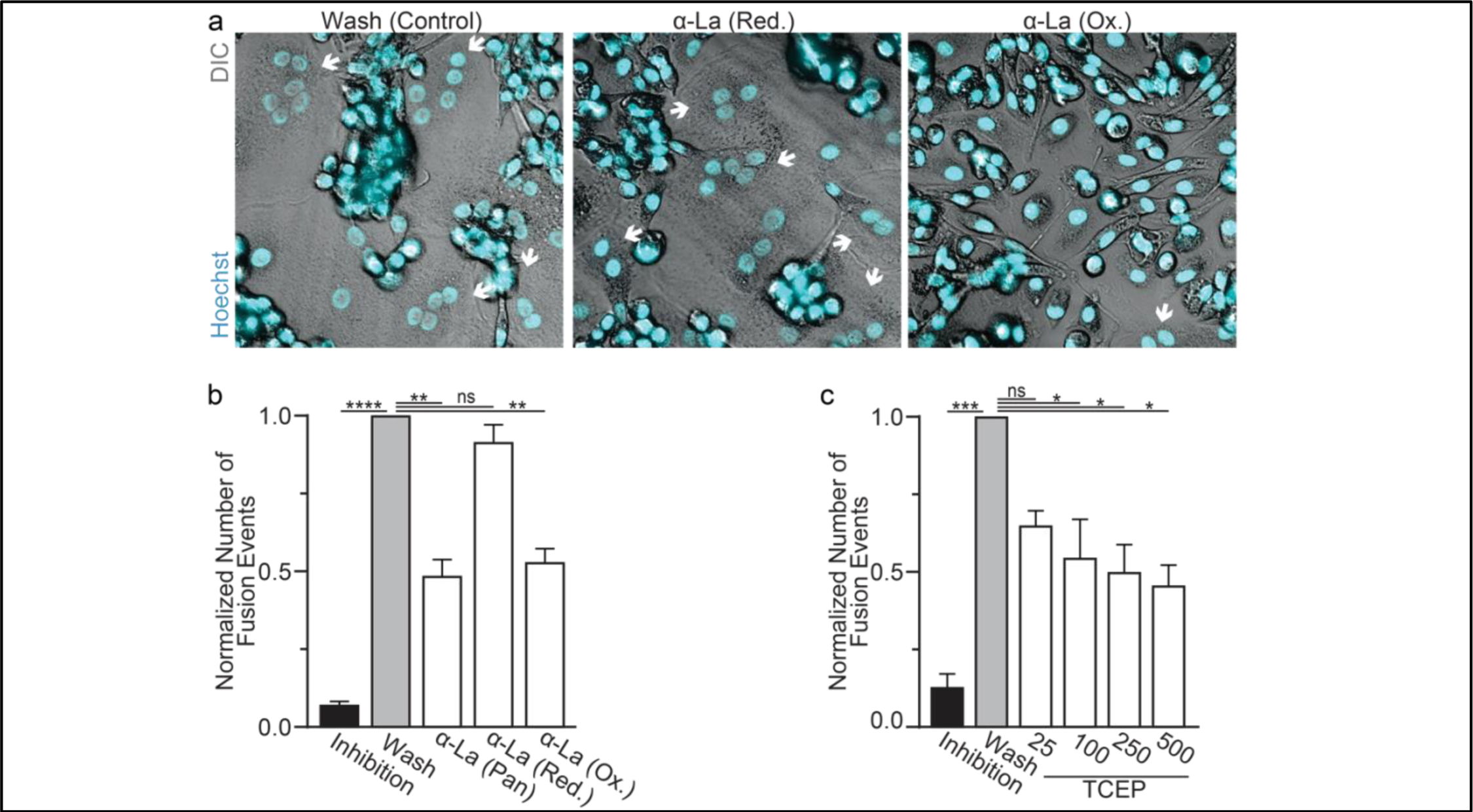
Oxidized La promotes osteoclast membrane fusion. **(a)** Representative fluorescence and DIC confocal micrographs of primary human osteoclasts following synchronized cell-cell fusion where hemifusion inhibitor was left (Inhibition), removed (Wash) or removed but the α-La antibodies indicated were simultaneously added. Cyan=Hoechst Arrows=Multinucleated Osteoclasts **(b)** Quantification of **a**. (n=5) (p= <0.0001, 0.0019, 0.44, and 0.0038, respectively) **(c)** Quantification of synchronized primary human osteoclast fusion events under control conditions or conditions where surface proteins are reduced (TCEP). Osteoclast fusion was synchronized by reversibly inhibiting cell-cell fusion using the membrane fusion inhibitor LPC. Inhibition=LPC applied and not removed, Wash=LPC applied and removed to allow synchronized fusion, TCEP=same as Wash with the addition of TCEP before LPC removal. (n=4, except 25 where n=2) (p= 0.0005, 0.36, 0.02, 0.016, and 0.013, respectively) Statistical significance evaluated via paired one-way ANOVA with Holm-Sidak correction.

To further validate the role of La’s redox status in osteoclast multinucleation and resorptive function, we chose to assess the functional impact of perturbing the oxidation status of recombinant La. Previously, we showed that the C-terminal half of La produced recombinantly and added to the medium bathing differentiating osteoclast precursors binds the surface of these cells and is sufficient to promote osteoclast fusion and resorptive function ^6^. We produced La 194-408 recombinantly and reduced a portion of the protein with TCEP followed by the application of iodoacetamide to block the free thiols in La 194-408, preventing their subsequent oxidation. When we compared the effects of La 194-408 vs reduced La 194-408 on primary human osteoclast multinucleation, we found that the reduction of recombinant La greatly diminished its ability to promote osteoclast multinucleation and resorption (Fig. 3a-c).

**Figure 3:**
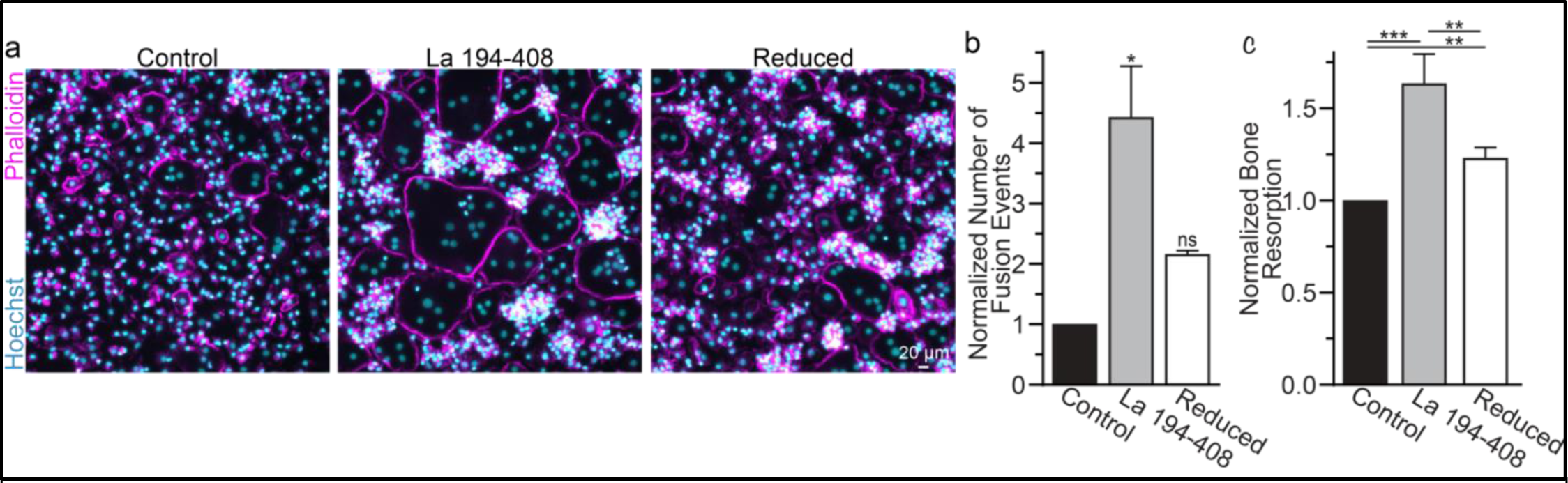
Surface La’s oxidation is functionally important for the formation and function of osteoclasts. **(a)** Representative fluorescence images of primary human osteoclast multinucleation following addition of control La 194-408 vs La 194-408 where cysteine residues were reduced by TCEP and then blocked by iodoacetamide treatments. Cyan=Hoechst. Magenta=Phalloidin **(b)** Quantification of osteoclast fusion events in **a**. (n=3) (p= 0.029 and 0.44, respectively) Statistical significance evaluated via paired Friedman test with Dunn’s correction. **(c)** Quantification of *in vitro* resorptive function in conditions described in **a**. (n=4) (p= 0.0001, .009 and 0.001, respectively) Statistical significance evaluated via paired one-way ANOVA with Holm-Sidak correction.

Many of the effects of redox signaling are mediated by the oxidation of cysteine residues within proteins ^40^. Human La has 3 cysteine residues. However, we have previously demonstrated that the N-terminal half of La (La 1-187), containing one of La’s cysteines, is dispensable for La’s role in promoting osteoclast fusion ^6^. Therefore, we focused on the two cysteines in the C-terminal half of La: Cys 232 and Cys 245. We recombinantly produced a La 194-408 mutant where cysteines 232 and 245 were mutated to alanine residues (La Cys Mut.) (Fig. 4a,b; Supp. Fig. 4a). Interestingly, we find that a fraction of La 194-408 migrates as a band ∼twice the predicted size of La 194-408 despite separation via denaturing gel electrophoresis, suggesting the presence of La 194-408 dimers^35,41^. In contrast, we observe no dimer band in La Cys Mut (Supp. Fig. 4 b,c). In addition, we find that the loss of Cys 232 and Cys 245 greatly diminishes the affinity of the α-La antibody that recognizes the oxidized La species, whereas both species are recognized by an α-6xHis monoclonal antibody via their N-terminal His tags (Fig. 4b). Finally, we found that the loss of Cys 232 and Cys 245 strongly abrogates the ability of La 194-408 to promote osteoclast fusion (Fig. 4c,d). From these data, we conclude that Cys 232 and Cys 245 – previously highlighted for their roles in redox-dependent structural changes in La ^35^ – are vital for La’s function in osteoclast multinucleation.

**Figure 4:**
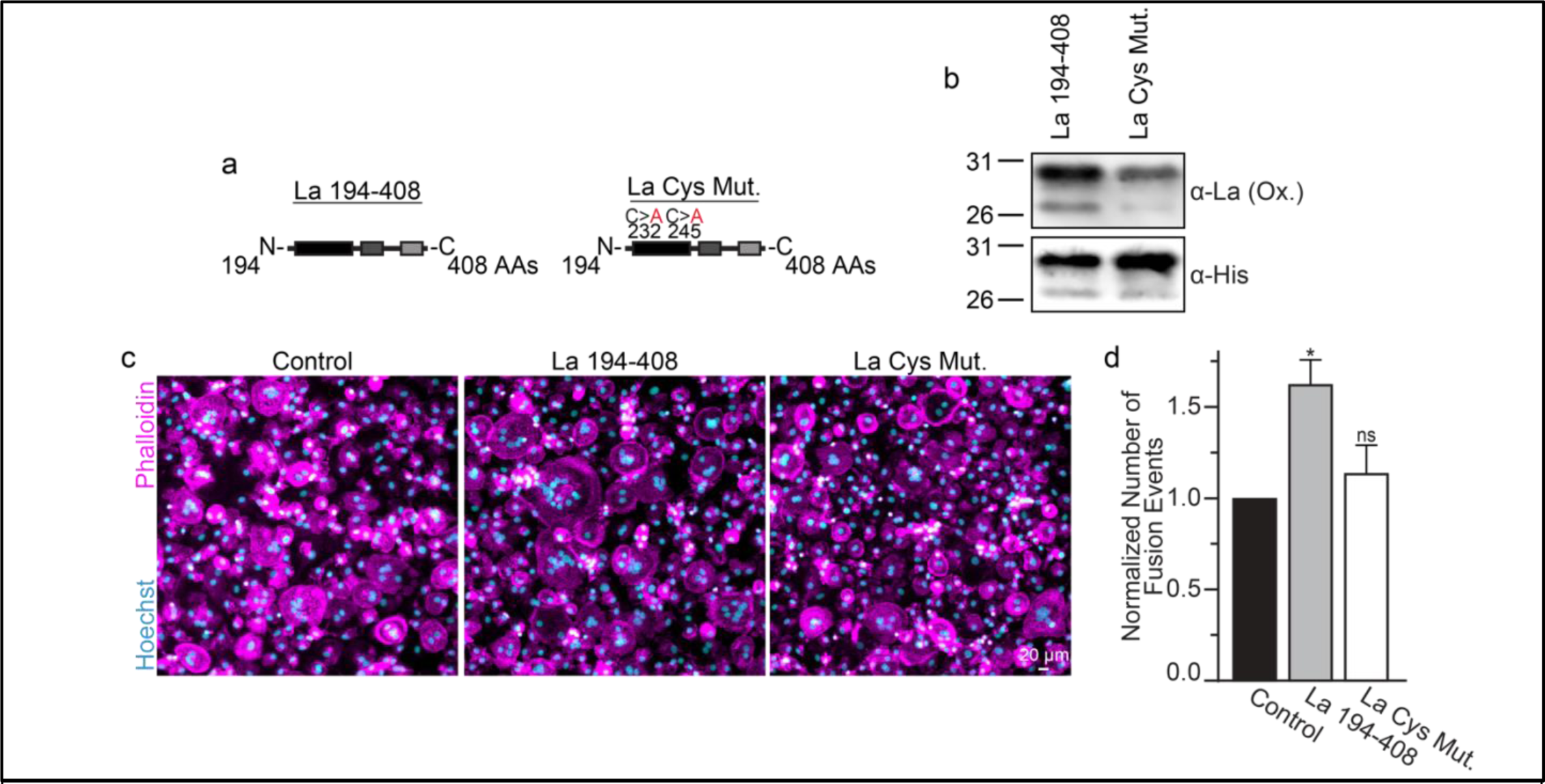
La 194-408 cysteine residues are critical for promoting osteoclast fusion. **(a)** Cartoons illustrating the domain structure of La’s C-terminal half and the location of its two cysteines. **(b)** Representative Western Blots depicting La C-terminal half and cystine mutant. **(c)** Representative immunofluorescence micrographs of fusing primary human osteoclasts under control conditions or treated with La **194-408** or cysteine mutant La **194-408**. **(d)** Quantification of the number of fusion events observed in c. (n=4) (p= 0.046 and 0.62, respectively) Statistical significance evaluated via paired one-way ANOVA with Dunnett correction.

To summarize, La promotes osteoclast fusion as an unconventional, cell surface associated, oxidized species. Moreover, our data suggest that the conformational transition from reduced to oxidized La species that is critical for the protein’s role in osteoclast formation and function depends on cysteine residues within its C-terminal half.

### La’s redox transition takes place in the cytoplasm and promotes surface delivery

The La pool within the nuclei of eukaryotic cells is primarily a reduced species ^35,36^. Finding that the surface La pool, which promotes multinucleation in osteoclasts, is enriched in an oxidized species raised the question of where La becomes oxidized? Is La oxidation in osteoclasts a consequence of surface delivery or does La oxidation precede surface trafficking?

Finding that La in permeabilized differentiating osteoclasts is recognized by the α-La antibody that recognizes oxidized rather than reduced La species (Fig. 1a and Supplement Fig. 1) suggested that the oxidation of La takes place in the cytoplasm of forming osteoclasts prior to delivery to the oxidizing environment of their surface. To further address this question, we utilized a cell-permeable reducing reagent N-Acetylcysteine (NAC) that has been widely used to inhibit intracellular ROS generation and suppress cytosolic redox signaling ^42,43^. Since under some conditions complex indirect effects of NAC raise rather than lower ROS levels ^44–46^, we verified the ROS-lowering effects of NAC in osteoclast precursors using a previously validated, cell-permeable ROS probe CellROX Deep Red ^24^. We found that NAC treatment suppresses ROS elevation stimulated by RANKL-initiated differentiation of human osteoclasts (Fig. 5a,b). While early treatment with NAC has been reported to disrupt the pre-fusion stages of osteoclast differentiation ^24^, we found that NAC treatment 24 hours after RANKL application (i.e., after early osteoclast precursor commitment), did not lower the steady state levels of osteoclastogenic differentiation factors (NFATc1, cFOS) or the transcripts of two fusion-related proteins (La or AnxA5) (Fig. 5c). We then explored the effects of suppressing ROS signaling on osteoclast formation and the trafficking of La, as in ^33,34^, we found that NAC mediated inhibition of ROS signaling suppresses osteoclast multinucleation in a dose dependent manner (Fig. 5d). Suppressing ROS signaling with NAC also inhibited the transition from reduced to oxidized, cytoplasmic La species, as evidenced by decrease in staining of permeabilized osteoclasts with α-oxidized La antibody and a complementary increase in staining of permeabilized osteoclasts with α-reduced La antibody (Supplement Fig. 5a-d). Most importantly, NAC application inhibited the cell-surface delivery of La in human osteoclasts (Fig. 5e). These findings suggest that NAC inhibits La function in osteoclasts by suppressing ROS-induced oxidation of La and its delivery to the surface of osteoclasts. Moreover, our findings also indicate that the transition from reduced to oxidized La species occurs inside the osteoclast cytoplasm during their commitment program rather than being a consequence of La arriving at the oxidizing environment of the extracellular milieu.

**Figure 5:**
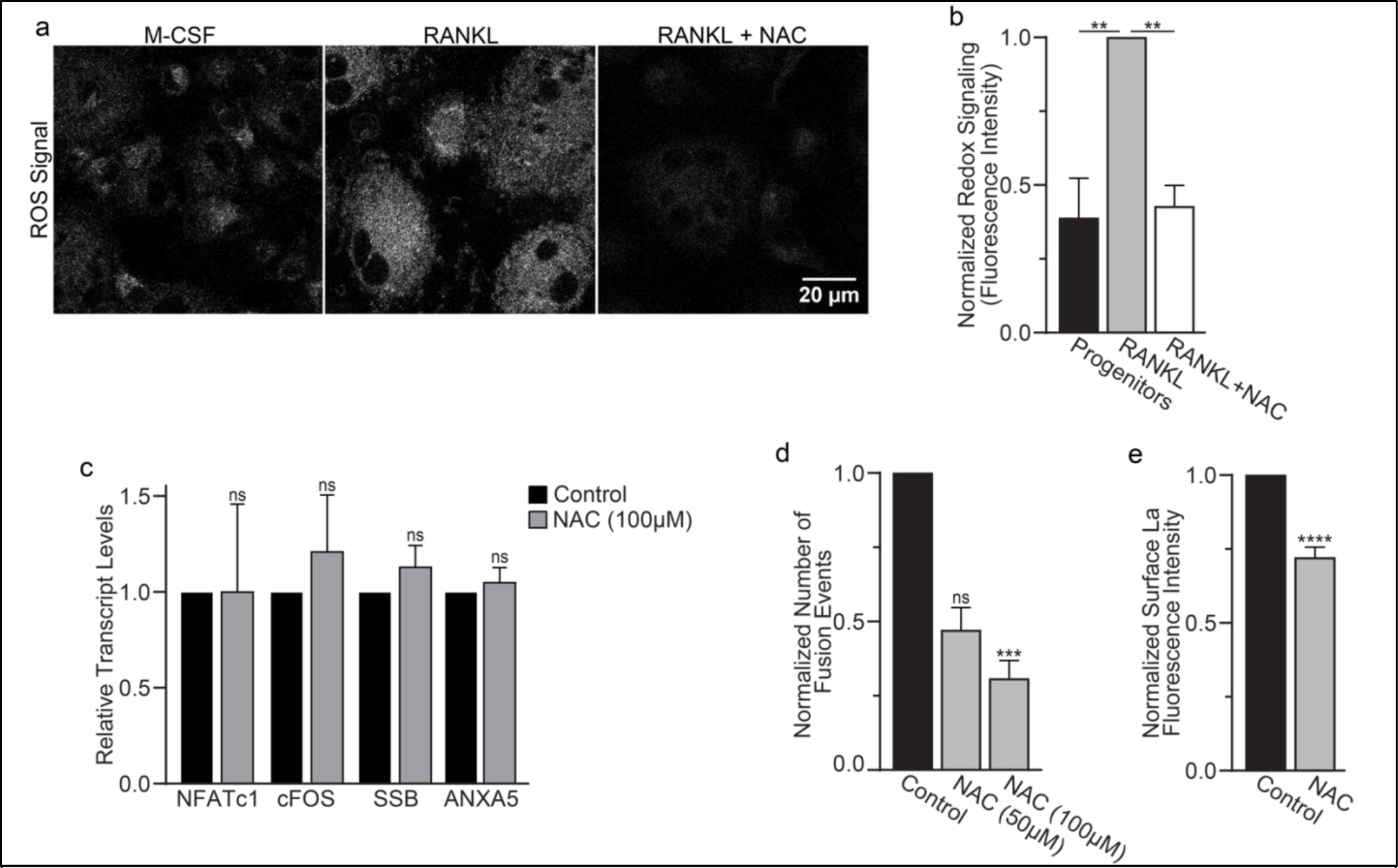
ROS promotes oxidized La surface trafficking and osteoclast fusion. **(a)** Representative confocal micrographs of ROS signal in primary human osteoclasts precursors under conditions lacking RANKL, following 16 hours of RANKL application, or following 16 hours and 1hr 100 μM NAC treatment. (Grey = CellRox Dye) **(b)** Quantification of ROS signaling in osteoclast progenitors, committed osteoclasts, or committed osteoclasts treated with the membrane permeable reducing reagent NAC (n=2, 4, and 3, respectively) (p= 0.004 and 0.0043, respectively) Statistical significance evaluated via paired one-way ANOVA with Holm-Sidak correction. **(c)** qPCR quantification of osteoclastogenesis markers (*NFATc1, cFOS*), La transcript (*SSB*), and annexin A5 (*ANXA5*). Expression evaluated in comparison to *GAPDH*. (n=4) (p= 0.50, 0.69, 0.32, and 0.45, respectively) Statistical significance evaluated via paired t-test. **(d)** Quantification of the number of fusion events observed between human osteoclasts in control conditions or conditions where fusion was inhibited via NAC treatment. (n=6) (p= 0.057 and 0.019, respectfully). Statistical significance evaluated via paired one-way ANOVA with Holm-Sidak correction. **(e)** Quantification of La surface staining of non-permeabilized cells with pan a-La antibodies at Day 3 post RANKL application without or with 50-100 μM NAC added at day 1 post RANKL application. (n=9) (p= <0.0001). Statistical significance evaluated via paired t-test.

While most of La in human cells is phosphorylated at Ser 366 ^47^, at the time of fusion, La is mostly dephosphorylated, as evidenced by a loss of the cell staining with antibodies specific for La phosphorylated at Ser366 (α-p366 La Ab) ^6^. We find that while little phosphorylated La is typically observed in the nuclei of differentiating osteoclasts during fusion timepoints, that NAC suppression of transient ROS signaling greatly increases phosphorylated La in osteoclasts (Fig. 6a,b). These data indicate that inhibition of intracellular ROS generation prevents La dephosphorylation and the loss of nuclear localization in differentiating osteoclast precursors. These findings substantiate the hypothesis that the redox signaling, which triggers a shift in La functional properties, is important for osteoclast fusion and takes place inside the cell.

**Figure 6.**
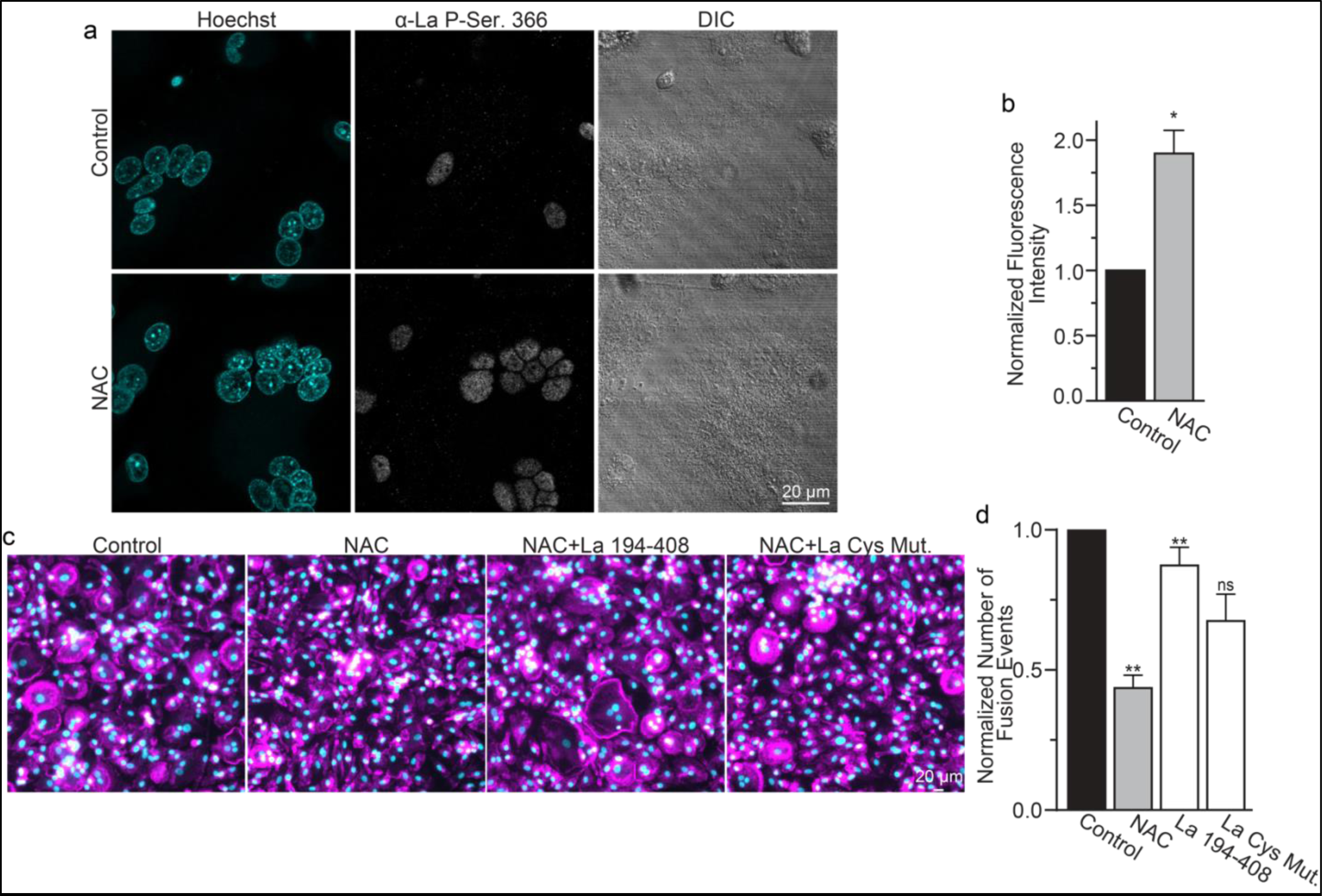
ROS signaling promotes dephosphorylation of La and osteoclast fusion by increasing the amounts of oxidized La at the surface of the cells. **(a)** Fluorescence microscopy and DIC images of permeabilized osteoclasts without or with application of 100 mm NAC (1h) stained with an α-La antibody that recognizes La phosphorylated at Ser366. **(b)** Quantification of the staining intensity from **a**. (n=3) (p=0.01). Statistical significance evaluated via paired t-test. **(c)** Representative fluorescence images of differentiating osteoclasts in control conditions, conditions where fusion was inhibited via NAC treatment (100 μM NAC added at day 2 days post RANKL application), and conditions where fusion was rescued by the application of recombinant La 194-408 or cysteine mutant La 194-408. **(d)** Quantification of the number of osteoclast fusion events in **c**. (n=5) (p= 0.001, 0.004, and 0.073, respectfully). Statistical significance were evaluated via One-way Anova with Holm-Sidak correction.

We found that NAC application both inhibits osteoclast multinucleation and suppresses delivery of oxidized La species to the cell surface of fusing osteoclasts (Fig. 5d,e). These findings motivated us to explore whether the suppressed osteoclast multinucleation caused by NAC treatment could be the result of deficient La delivery to the cell surface. If the suppressed fusion observed following NAC treatment is a consequence of suppressing a ROS triggered “La trafficking switch”, then perhaps simply adding La to the medium bathing NAC-treated cells can rescue La surface pools and fusion? Indeed, we found that application of La 194-408 to NAC-treated cells rescued osteoclast fusion inhibition (Fig. 6c,d). However, the ability of La 194-408 to rescue NAC-inhibited osteoclast fusion is at least partially dependent on Cys 232 and Cys 245, as mutation of these residues to Ala greatly diminished the ability of La 194-408 to rescue NAC effects on fusion. Finding that the NAC-mediated ROS suppression of surface trafficking and fusion can be compensated for by the application of exogenous La strongly supports the conclusion that ROS signaling plays a vital role in triggering La’s delivery to the surface of osteoclasts and the promotion of their multinucleation. Moreover, the loss of La’s ability to rescue NAC suppression of osteoclast multinucleation when Cys 232 and Cys 245 are mutated to Ala also suggests that these residues are vital for oxidized La’s ability to promote osteoclast multinucleation and resorptive function.

In summary, ROS signaling downstream of osteoclast commitment contributes to the cell fusion stage of osteoclast formation by promoting intracellular oxidation of La and its delivery to the surface of fusion-committed cells, where La promotes multinucleation and subsequent resorptive activity in human osteoclasts (Fig. 7).

**Figure 7:**
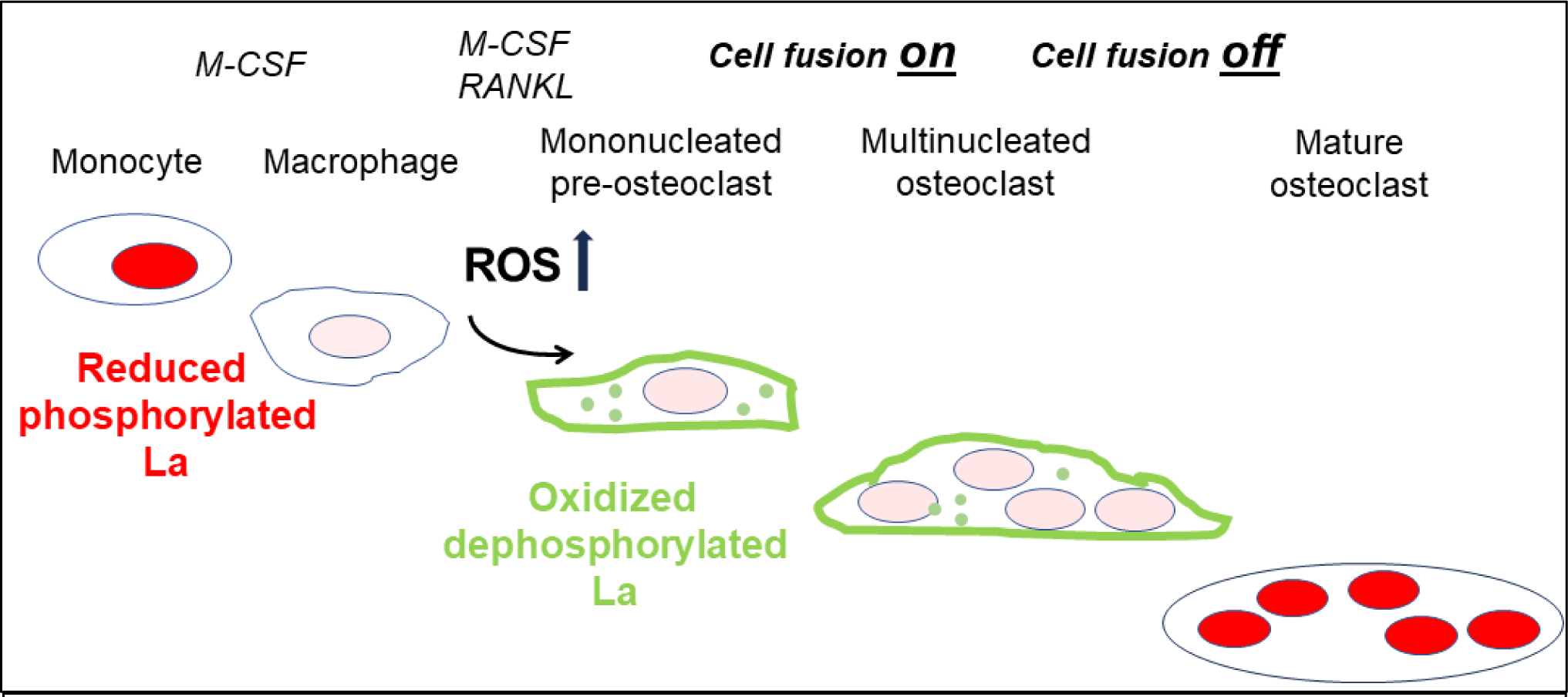
ROS signaling induced restructuring of La from reduced to oxidized species triggers La re-localization from nucleus to the surface of differentiating osteoclasts and promotes their fusion and resorptive function. An illustrated depiction of osteoclastogenic differentiation from monocytes to mature bone-resorbing osteoclasts. Machrophage precursors are derived via the M-CSF activation of circulating monocytes. Osteoclastst differentiation is initiated by subsequent application of M-CSF and RANKL, which ellicits intracellular ROS production leading to drastic changes in the redox state and localization of La. La transitions from a prodominatly nuclear, reduced species of La in monocytes and macrophages to an oxidized, dephosphorylated species that traffics to and associates with the surface of fusion-competent osteoclasts. When osteoclast arrive at an appropriate size and fusion stops, thee mature multinucleated osteoclasts exhibit a predominately nuclear, reduced La species, as is typical of other eukaryotic cells.

## DISCUSSION

The reversible shift of redox homeostasis in cells to a moderately oxidized state, referred to as redox signaling, regulates the function and localization of many proteins and favors differentiation vs. proliferation ^48–50^. Our findings confirm earlier reports emphasizing the importance of redox signaling in osteoclastogenic differentiation ^17,23,24^. ROS signaling and its inhibition by NAC likely influence osteoclast formation in many ways and at many points in the formation of multinucleated osteoclasts ^24^. However, our finding that exogenous La rescues the NAC-mediated suppression of osteoclast fusion supports the hypothesis that ROS signaling promotes cell fusion stage of osteoclastogenesis by triggering La’s delivery to the surface of osteoclasts. This mechanism enriches our recent identification of La protein as a regulator of the cell-cell fusion stage of osteoclast formation ^6^ and highlights the role of osteoclast ROS signaling in directing fusion machinery to the surface of osteoclast progenitors. While the reduced species of La functions as an essential nuclear RNA chaperone, oxidized La shifts to the cytoplasm and shuttles to the surface of the osteoclasts. It is this specialized oxidize species of cell-surface associated La that promotes osteoclast fusion and bone resorption.

The transition from reduced to oxidized species of La is accompanied by many changes in the protein structure, including an oxidation-induced oligomerization of the protein, a loss of almost half of its helical content, and a change in accessibility of epitopes recognized by conformation specific antibodies ^35^. Our data show that La belongs to a category of fold switching proteins that change functions in response to the cellular environment, more specifically, oxidizing and reducing intracellular conditions ^51^. In addition to direct changes in the structure of La caused by its oxidation, ROS can influence La function in osteoclast fusion by indirect effects. Formation of multinucleated osteoclasts involves the caspase 3-cleaved, non-phosphorylated species of La ^6^ and ROS can influence La properties by activation of both caspase 3 ^52^ and PP2A-like phosphatase ^53^, previously reported to dephosphorylate La Ser366 before La cleavage ^54^. Indeed, while our finding that NAC inhibits dephosphorylation of La can indicate that redox-dependent changes in La’s conformation are required for the dephosphorylation of the protein, it can also be explained by the ROS dependence of PP2A activity. In both scenarios, redox signaling promotes intracellular La modifications and trafficking that delivers the fusion promoting species of La to the surface of cells.

We still do not know how cell-surface La promotes osteoclast fusion and why the oxidized species of La is functional in this unique cellular context. For many viral and intracellular fusion processes, remodeling of membrane bilayers in fusion is thought to be driven by the conformational energy released at the time and place of fusion in the restructuring of fusion-promoting proteins ^55^. Finding that La acquires an oxidized conformation already in cytoplasm rather than at the cell surface at the time of fusion argues against the hypothesis that redox restructuring of La directly contributes to membrane remodeling. Furthermore, our finding that La association with the cell surface does not change after reducing oxidized La with TCEP argues against the hypothesis that La oxidation is merely required for its association with the surface of osteoclasts.

Our data, which indicate that fusion competence in osteoclast precursors depends on ROS signaling, can be, at least partially, explained by the oxidation-dependent changes in the structure of La protein and its nucleo-cytoplasmic-cell surface shuttling. Formation of disulfide bonds; re-localization of normally nuclear proteins to the cytoplasm, extracellular medium, and cell surface; and dramatic changes in function in response to transient increases in ROS concentrations have been well described for other nuclear chaperones, in particular, high mobility group box 1 (HMGB1) ^19,56,57^. Like La, HMGB1 has three cysteines and mild HMGB1 oxidation generates disulfide bonds that stabilize homodimers^58^. Homodimerization by formation of disulfide bonds is also a pre-requisite for the unconventional secretion of Fibroblast Growth Factor 1 ^59^.

The specific pathway(s), by which RANKL-induced increases in ROS levels ^24^ shift the redox state of La and facilitate its unconventional secretion, remain to be clarified. Moreover, we hope to explore the contributions of related pathways like the production of nitric oxide (NO) and other reactive nitrogen species. In the case of HMGB1, the secretion of the protein, allowing it to act as a signaling molecule outside the cell, in addition to ROS, depends on NO signaling mediated by NO binding of one of a cysteine thiol in protein ^57^. Interestingly, like redox signaling ^17,23,24^, NO signaling promotes osteoclastogenesis ^60^ and the translocation of La protein to the cytoplasm ^36^. More work is needed to explore the contributions of NO signaling to La function in osteoclasts and to compare the molecular mechanisms that deliver these two redox-, fold-, and function-shifting proteins, La and HMGB1, to the cell surface and extracellular medium.

In conclusion, in this study, we identified redox signaling as a molecular switch that redirects La protein away from the nucleus, where it protects precursor tRNAs from exonuclease digestion, and towards its separable function at the osteoclast surface, where La regulates the multinucleation and resorptive functions of these managers of the skeleton.

Proteins involved in osteoclastogenesis represent potential therapeutic targets for treating bone loss diseases. Finding that osteoclast La promotes bone-resorption by acting not only at a different location but also in a different conformation from in comparison to the species of La that carries out its essential and ubiquitous RNA chaperoning functions (cell surface vs. nucleus and oxidized vs. reduced form), may help in minimizing off target effects of La targeting treatments.

## Acknowledgements

We thank the National Institutes of Health Department of Transfusion Medicine for isolating the monocytes used in this study. Our work was supported by the Intramural Research Program of the Eunice Kennedy Shriver National Institute of Child Health and Human Development, National Institutes of Health and by the of Research on Women’s Health (ORWH) through the Bench to Bedside Program award # 884515.

## METHODS

### Reagents

Human M-CSF and RANKL were purchased from Cell Sciences (catalogue #CRM146B and #CRR100B, respectively). LPC (1-lauroyl-2-hydroxy-sn-glycero-3-phosphocholine, #855475); PC (1,2-dioleoyl-sn-glycero-3-phosphocholine, #850375 C) was purchased from Avanti Polar Lipids. Bone Resorption Assay Kits were purchased from Cosmo Bio Co. (Catalogue # CSR-BRA-24KIT) and used according to the manufacturer’s instructions. Hoechst 33342 and phalloidin-Alexa555 were purchased from Invitrogen (#H3570 and A30106, respectively). TCEP (Tris (2-carboxyethyl) phosphine) was purchased from Thermo Fisher Scientific (Pierce™ TCEP-HCl; catalogue # A35349). NAC (N-Acetyl-L-cysteine) and sodium iodoacetamide were purchased from Sigma Aldrich (catalogue # A9165 and GERPN6302).

### Cells

Elutriated monocytes from healthy donors were obtained through the Department of Transfusion Medicine at National Institutes of Health under protocol 99-CC-0168 approved by the National Institutes of Health Institutional Review Board. Research blood donors provided written informed consent and blood samples were de-identified prior to distribution, Clinical Trials Number: NCT00001846. We also used elutriated monocytes from healthy donors obtained through Elutriation Core Facility, University of Nebraska Medical Center, informed consent was obtained under an Institutional Review Board approved protocol for human subject research 0417-22-FB. Research blood donors provided informed consent and samples were de-identified prior to distribution. Primary human osteoclasts were derived as described previously ^6^. Briefly, elutriated monocytes were added to complete media [α-MEM (Gibco) +10% FBS (Gibco)+ 1x penicillin/streptomycin/glutamate (Gibco)] supplemented with 100 ng/ml recombinant M-CSF and plated at 1×10^6^ cells/ml for 6 days (refreshing media at Day 3). Next, the cells were placed into complete medium supplemented with 100ng/ml recombinant M-CSF and 100 ng/ml recombinant RANKL to induce osteoclastogenesis 3-4 days to obtain multinucleated, resorption competent human osteoclasts.

### Antibodies

Murine monoclonal α-La antibodies that specifically recognize oxidized and reduced species of La or both species of La (7B6, 312B and 5B9, respectively, ^35^, referred to as oxidized La Ab, reduced La Ab and pan α-La Ab) were described and characterized by the laboratory of Dr. Michael Bachmann. The antibodies were produced recombinantly as described in ^35^.

We also used rabbit anti-La Phospho-Ser366 antibody (Abcam, 61800), referred to as α-p366 La Ab that recognizes phosphorylated human La (phosphoSer366). We also used an additional monoclonal murine α-La antibody (α-La mAb; Abcam, catalogue #75927) that we found to recognize both oxidized and reduced forms of La. A α-6xhis murine monoclonal antibody (Abcam, ab18184) was used to recognize the 6xhis tag covalently modifying our recombinantly produced La protein fragments and α-Cyclophilin B (Cell Signaling Technology, D1VdJ Rabbit mAb #43603) as a loading control.

### Constructs & Recombinant Protein

Constructs encoding recombinant La 194-408 and the cysteine mutants used in this manuscript were previously described, characterized, and provided by the Bachmann Lab. Each were transformed into BL2 (DE3) chemically competent *E. coli* (Thermo Fisher Scientific) and recombinant protein production was induced via IPTG induction (Sigma). Cells were lysed with BugBuster HT (Millipore) supplemented with protease inhibitors (Complete, Pierce), 6xHis-La proteins were purified using HisPur Cobalt Spin columns (Thermo Fisher Scientific), and endotoxin was removed via Pierce high-capacity endotoxin removal columns (Thermo Fisher Scientific), each according to manufacturer’s instructions. Proteins were sterile filtered, aliquoted, and kept at −80 °C. Some La 194-408 was irreversibly modified via 45 min incubation with 10mM TCEP followed by a 45 min incubation with 10mM iodoacetamide. Modified protein was then subsequently exchanged into PBS (Gibco) using 10K MW concentrators according to manufacturer’s instructions (Amicon). Control La 194-408 for these experiments was treated identically, except for the omission of TCEP and iodoacetamide.

### Microscopy

For high-resolution immunofluorescence analysis of protein localization, we washed cells with PBS and then rapidly fixed with warm, freshly prepared 4% formaldehyde in PBS at 37 °C. The cells were subsequently washed with PBS. To permeabilize cells, we incubated them for 5 min in 0.1% Triton X100 in PBS. The cells were subsequently stained in PBS supplemented with 10% FBS for 10 min at room temperature to suppress non-specific binding. Then, cells were incubated with primary antibodies for 1 hr in PBS supplemented with 10% FBS. After 5 washes in PBS, we incubated the cells with fluorescent secondary FAB fragments raised to the species corresponding to the primary antibody for 1 hr in PBS supplemented with 10% FBS (either Anti-rabbit IgG Fab2 Alexa Fluor ® 555 or Anti-mouse IgG Fab2 Alexa Flour ® 488, both Cell Signaling Technology, Catalogue # 647 4414 S and # 4408 S, respectively, in 1:500 dilution) and then washed 5 times with PBS prior to imaging. For non-permeabilized conditions, we followed the same protocol, but omitted all use of detergents.

Images were captured on a Zeiss LSM 800, confocal microscope using a C-Apochromat 63x/1.2 water immersion objective lens.

For basic immunofluorescence analysis of cell fusion and morphology, we stained osteoclast cytoskeletal boundaries with Phallodin-Alexa Flour ® 488 (Thermo Fisher Scientific, 1:2000) and nuclei with Hoechst 33342 (Thermo Fisher Scientific, 1:5000). Cells were fixed as described above, washed with PBS, and stained with toxin/dyes for 1 hr in complete staining buffer (PBS + 5% FBS + 0.1% TX100) before a final PBS wash prior to imaging. 10 selected fields of view were imaged on a grid from the center of each well/dish in automated fashion using Alexa488, Hoechst and phase contrast compatible filter cubes (BioTek) on a Lionheart FX microscope using a 10x/0.3 NA Plan Fluorite WD objective lense (BioTek) using Gen3.10 software (BioTek). Each image was separated by approximately 1800 microns in both x and y parameters.

All image data were evaluated using Fiji/ImageJ’s open-source image processing package v.2.1.0/1.53c.

### Cell Fusion Quantitation

Osteoclast fusion efficiency was evaluated as the number of fusion events between cells in 10 images. In brief, since regardless of the sequence of fusion events, the number of cell-to-cell fusion events required to generate syncytium with N nuclei is always equal to N-1, we calculated the fusion number index as Σ (Ni − 1) = Ntotal − Nsyn, where Ni = the number of nuclei in individual syncytia and Nsyn = the total number of syncytia. We normalized the number of fusion events to the total number of nuclei (including unfused cells) to control for small variations in cell density from dish to dish and image to image. In contrast to traditional fusion index measurements, this approach gives equal consideration to fusion between two mononucleated cells, one mononucleated cell and one multinucleated cell and two multinucleated cells. In traditional fusion index calculations, fusion between two multinucleated cells does not change the percentage of nuclei in syncytia. If instead one counts the number of syncytia, a fusion event between two multinucleated is not just missed but decreases the number of syncytia. In contrast, the fusion number index is inclusive of all fusion events.

### Synchronization of osteoclast fusion

Osteoclast fusion was synchronized as described in ^38^. Briefly, osteoclast media was refreshed with media supplemented with 100 ng/ml M-CSF, 100 ng/ml RANKL and 350 μM lauroyl-LPC 72 h post RANKL treatment. Following 16 h, LPC was removed via 5 washes with fresh media and cells were allowed to fuse in the presence or absence of antibody treatment or recombinant La at the concentrations described in the figure legends for 90 min.

### Transcript Analysis

For real-time PCR, total RNA was collected from cell lysates using PureLink RNA kit following the manufacturer’s instructions (Invitrogen # 12183018 A). cDNA was generated from total RNA via reverse transcription reactions using a High-Capacity RNA-to-cDNA kit according to the manufacturer’s instructions (Applied Biosystems, # 4387406). cDNA was then amplified using the iQ SYBR Green Supermix (Biorad). All primers were predesigned KiCqStart SYBR Green primers with the highest rank score specific for the gene of interest or GAPDH control and were used according to the manufacturer’s instructions (Sigma). All Real-time PCR reactions were performed and analyzed on a CFX96 real-time system (Biorad), using GAPDH as an internal control. Fold-change of gene expression was determined using the ΔΔCt method. 3–4 independent experiments were performed, and each was analyzed in duplicate.

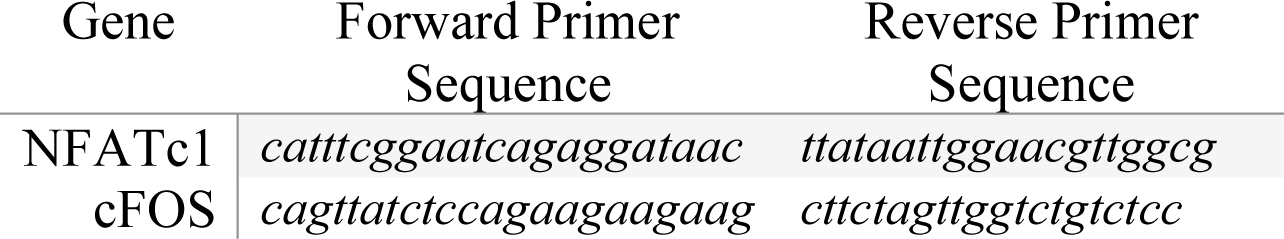
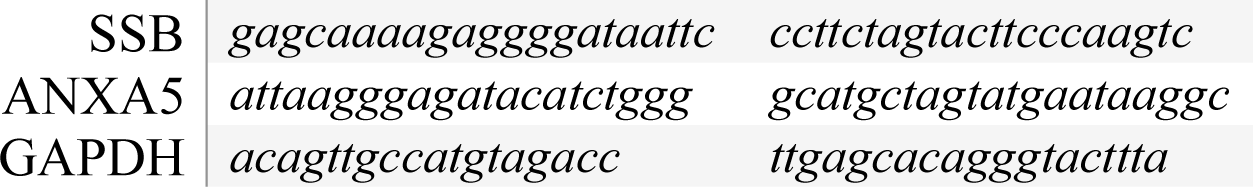

### Mineral Resorption

Mineral resorption was evaluated using mineral resorption assay kits from Cosmo Bio USA according to the manufacturer’s instructions. In short, fluoresceinamine-labeled chondroitin sulfate was used to label 24-well, calcium phosphate-coated plates. Human, monocyte-derived osteoclasts were differentiated as described above, using alpha MEM without phenol red. Media were collected at 4–5 days post RANKL addition, and fluorescence intensity within the media was evaluated as recommended by the manufacturer. Data were normalized to the level of fluorescence released by control cells where RANKL was not added.

### Statistical Analysis

Statistical analyses were performed using Prism software (GraphPad Prism version 8.0.0). Unless stated in the legend, differences between groups were observed in each experiment, cells from each donor were paired across the conditions described, and statistical significance was assessed via Student’s t test. Due to the inherent variability in the derivation of primary human monocytes to osteoclast, we analyzed statistical significance using a ratio paired t test, where the raw values for the assay are logarithmically transformed and then assessed, when the precise time course of osteoclast differentiation and baseline extents of fusion varied considerably from donor to donor. The quantitated results presented all represent the mean ± the standard error of the mean (SEM). While the p values for each statistical comparison are defined in the legends of each figure, we graphically represented our statistical evaluations using the following symbols: ns, p= >0.05; *, p= <0.05; **, p=<0.01; ***, p=<0.001.

---

**Supplementary Figure 1.**
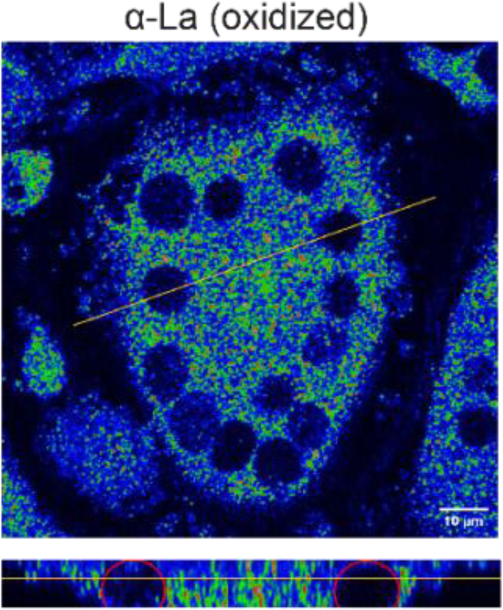
Oxidized La is found in the cytosol. 3D stack image depicting the cytosolic localization of oxidized LA. Top - XY-slice slightly below the equatorial plane of a permeabilized multinucleated osteoclast (3 days post-RANKL application) stained with an α-La antibody that recognizes oxidized La. 3D stack of this representative cell was acquired with 0.22μm step interval. Bottom - Z-slice through the orange line on the top panel. Red ellipses show the approximate outlines of two nuclei in the slice, and the orange line shows the location of the XY-slice shown on top.

**Supplementary Figure 4:**
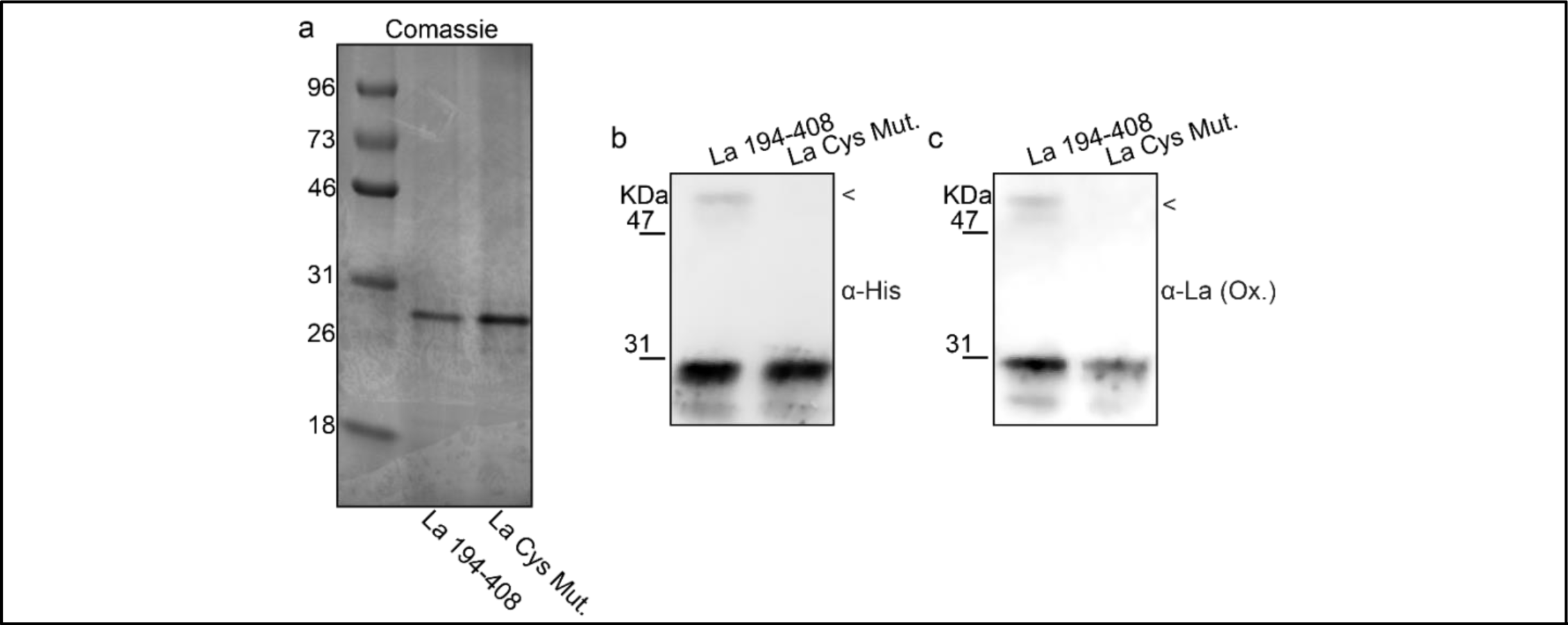
La C-terminal half and cysteine mutant purification. **(a)** A gray-scale image of La 194-408 or cysteine mutant La 194-408 separated via polyacrylamide gel electrophoresis and visualized using Coomassie staining. Representative Western Blots depicting La C-terminal half and cysteine mutant recognized by α-6xhis **(b)** or α-La (ox.) α-6xhis **(c)**. < Denotes the migration of La 194-408 as a dimer.

**Supplemental Figure 5.**
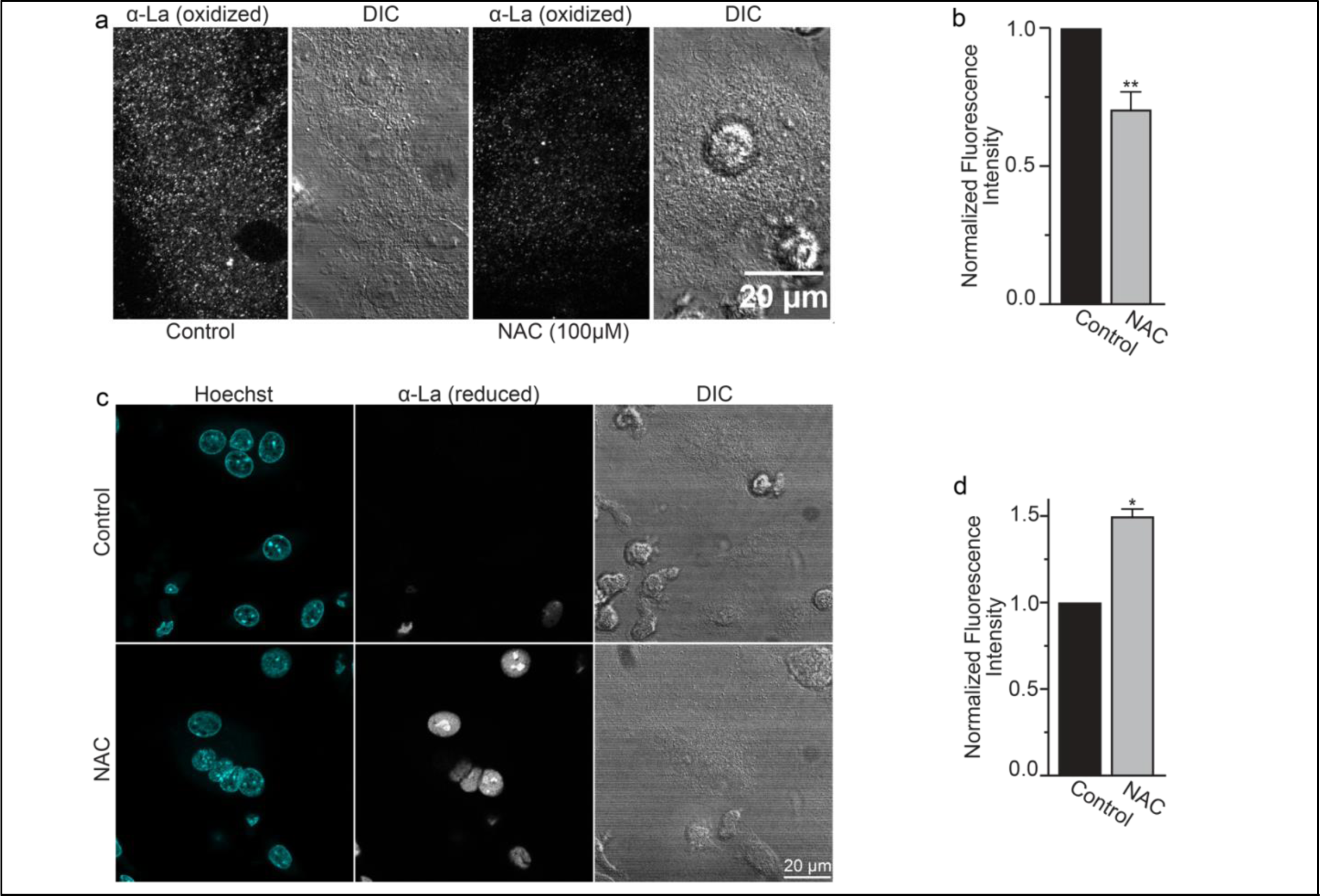
NAC inhibits redox shift from reduced to oxidized species of intracellular La. **(a)** Fluorescence microscopy and DIC images of permeabilized osteoclasts treated or not treated (control) with 100 mm (1h) NAC and stained with an α-La antibody that recognizes oxidized La at 3 days post RANKL application. **(b)** Quantification of **a**. (n=5) (p=0.009). **(c)** Fluorescence microscopy and DIC images of permeabilized osteoclasts treated or not treated (control) with 100 mm (1h) NAC and stained with an α-La antibody that recognizes reduced La at 3 days post RANKL application. **(d)** Quantification of **c**. (n=2) (p=0.04). Statistical significance evaluated via paired t-test.

